# Connecting the dots: applying multispecies connectivity in marine park network planning

**DOI:** 10.1101/2023.11.22.568386

**Authors:** Katie Gates, Jonathan Sandoval-Castillo, Andrea Barceló, Andrea Bertram, Eleanor A. L. Pratt, Peter R. Teske, Luciana Möller, Luciano B. Beheregaray

**Affiliations:** Molecular Ecology Lab, College of Science and Engineering, Flinders University, GPO Box 2100, Adelaide, SA 5001, Australia; Cetacean Ecology, Behaviour and Evolution Lab, College of Science and Engineering, Flinders University, GPO Box 2100, Adelaide, SA 5001, Australia; Centre for Ecological Genomics and Wildlife Conservation, Department of Zoology, University of Johannesburg, Auckland Park 2006, South Africa

**Keywords:** marine protected areas, gene flow, connectivity networks, spatial conservation prioritisation, seascape genomics

## Abstract

Marine ecosystems are highly dynamic, and their connectivity is affected by a complex range of biological, spatial, and oceanographic factors. Incorporating connectivity as a factor in the planning and management of marine protected areas (MPAs) is important yet challenging. Here, we used intraspecific genetic and genomic data for five marine species with varying life histories to characterise connectivity across a recently established South Australian MPA network. We generated connectivity networks, estimated cross-species concordance of connectivity patterns, and tested the impact of key spatial and oceanographic factors on each species. Connectivity patterns varied markedly among species, but were most correlated among those with similar dispersal strategies. Ordination analyses revealed significant associations with both waterway distances and oceanographic advection models. Notably, waterway distances provided better predictive power in all-species combined analyses. We extended the practical relevance of our findings by employing spatial prioritisation with Marxan, using node values derived from both genetic and geographic connectivity networks. This allowed the identification of several priority areas for conservation, and substantiated the initial decision to employ spatial distance as a proxy for biological connectivity for the design of the South Australian marine park network. Our study establishes a baseline for connectivity monitoring in South Australian MPAs, and provides guidelines for adapting this framework to other protected networks with intraspecies genetic data.

## 1. Introduction

Ecosystem connectivity is an integral aspect of planning design in protected areas networks, as it affects the structure, function, and dynamics of populations and communities (Carr et al. 2003, Carr et al. 2017, Grummer et al. 2019). One important component of this is population connectivity, i.e., the dispersal of individuals between spatially separated populations (Treml et al. 2008, Cowen and Sponaugle 2009). Population connectivity is highly relevant to both biodiversity conservation and fisheries management because it directly impacts demographic processes such as colonisation, recruitment, growth, and decline (Hastings and Botsford 2006, Aiken and Navarrete 2011). It also affects the distribution of genetic diversity, which can influence a population’s evolutionary viability and adaptive resilience to changing environments (Hoffmann and Sgro 2011, Frankham et al. 2017). In the marine environment, the design of protected areas does not often include explicit criteria for connectivity; instead, it is frequently based on surrogate measures such as geographic distance (Leslie 2005, DEH 2008). This may overlook other important modulators such as ocean circulation, distribution of traversable habitat, organisms’ life histories and population sizes, and temporal fluctuations of any of these factors (Weeks 2017, Balbar and Metaxas 2019). Fortunately, biologically informed data are becoming increasingly available to researchers, improving prospects for data-driven decision making in spatial conservation prioritisation.

Genetic and genomic tools are particularly valuable for population connectivity assessments. Rather than relying on individual and potentially stochastic dispersal events, as for example in tagging or photographic recapture studies, genetic information has the advantage of capturing patterns of both short- and long-term demographic exchange. Moreover, since population genetic approaches provide a measurement of realised connectivity, they can be used to ground-truth predictive models for dispersal based on physical ocean circulation (Bracco et al. 2019, Wilcox et al. 2023). Ocean currents facilitate the passive dispersal of many marine species (Cowen and Sponaugle 2009), and are also likely to affect population structuring of active dispersers due to influences on local habitats and their prey (Möller et al. 2007, Hays 2017). Importantly, oceanographic models, when combined with genetic data, might allow forecasting of advective connectivity changes under future climate scenarios (Coleman et al. 2017), or project the flow of climate-adaptive genetic variation into more vulnerable populations (Boulanger et al. 2020). Successful integration of advection models and genetic approaches could therefore pre-emptively inform conservation prioritisation to maximise the resilience of marine species in a warming climate.

Population genetic and genomic datasets are rapidly accumulating for a variety of marine species and seascapes around the world (Riginos et al. 2016; Grummer et al. 2019). Likewise, genetic connectivity and population structure have been studied in a growing number of Australian marine taxa (reviewed by Riginos et al. 2016, Teske et al. 2017, Jones et al. 2018). However, such studies have typically focussed on single species, limiting inferences of broader ecosystem management (Jones et al. 2018). Yet, academic publishers are moving gradually towards open data policies, and if data can be integrated from multiple independent studies, there is an opportunity to characterise regional patterns of connectivity more comprehensively.

In this study, we took a meta-analytical approach to assess the connectivity of multiple species across the South Australian Representative System of Marine Protected Areas (SARSMPA). The SARSMPA is a network of 19 multiple-use marine parks distributed across South Australia’s bioregions that was established in 2009 (Figure 1). Our meta-analysis was motivated by the need to address key evaluation questions in the Marine Parks Monitoring, Evaluation and Reporting Program (Bryars et al. 2017). These were: “to what extent have marine parks strategies contributed to the maintenance of ecological processes?” and “to what extent have marine parks strategies contributed to enabling marine environments to adapt to impacts of climate change?”. These key knowledge gaps are expected to be encountered during the design and spatial monitoring of marine parks around the world, attesting to the broader relevance of our study. We re-analysed existing genetic and genomic datasets in an integrative framework to provide information about connectivity patterns across the SARSMPA and the relative variation among life history types, including both active and passive dispersers. We evaluated relationships between multispecies connectivity and hypothesised environmental influences, including waterway distances and simulated estimates of advection connectivity. Finally, we applied these results to identify priority conservation areas for maintaining intraspecific genetic connectivity, and outline opportunities to extend our framework under alternative sampling designs. Our integrated analytical framework provides an exemplar of applying multispecies connectivity planning that can be readily adapted to marine protected networks elsewhere in the world as long as suitable intraspecies genetic or genomic data are available. The continued integration of biological and physical data will be invaluable in improving spatial conservation planning and monitoring of marine ecosystems, especially in the face of climate change and other anthropogenic pressures.

**Figure 1.**
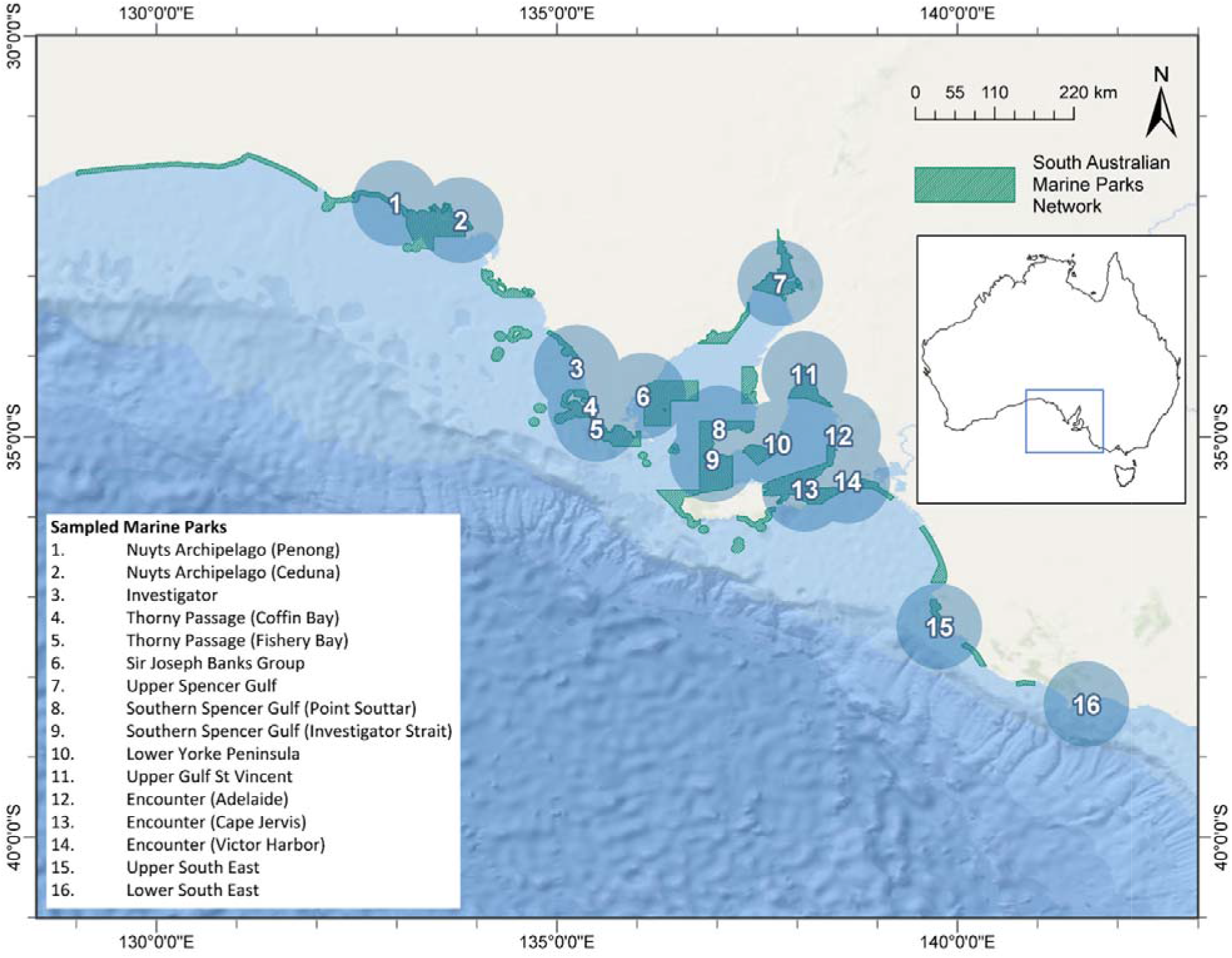
Sampling localities across the South Australian Marine Parks network based on aggregated data from the five study species, *Delphinus delphis*, *Tursiops aduncus*, *Chrysophrys auratus*, *Siphonaria diemenensis*, and *Nerita atramentosa*. Inset shows the study region in the map of Australia.

## 2. Methods

### 2.1 Cataloguing and integrating existing genetic and genomic datasets

We selected available genetic and genomic datasets for which species’ ranges and respective sampling schemes covered a broad region of the SARSMPA, with >3 South Australian sampling localities. This was further narrowed down to include only the two most common marker types used in population genetic analyses, microsatellites, and single nucleotide polymorphisms (SNPs). While SNPs generally have greater precision because they originate from a large number of loci throughout the genome, high concordance is expected for estimates of genetic diversity and population structure between these marker types (Zimmerman et al. 2020). For microsatellites, we used the full datasets described in the original publications (Teske et al. 2015, 2016, Teske et al. 2017). Unlike microsatellites, which are assumed to be selectively neutral, SNPs can be neutral or adaptive. Given that variation under local environmental selection may bias demographic inferences, we used datasets from which putatively adaptive variants had already been removed (as described in Barceló et al. 2021, Bertram et al. 2022, Pratt et al. 2022, Bertram et al. 2023). Based on these requirements, we identified suitable high-quality datasets for five species (Figure 2, Table 1). These included two iconic and legally protected cetaceans (the common dolphin, *Delphinus delphis*, and the Indo-Pacific bottlenose dolphin, *Tursiops aduncus*), a commercially, culturally, and recreationally valuable teleost (Australasian snapper, *Chrysophrys auratus*), and two ecologically important intertidal invertebrates (Van Diemen’s siphon limpet, *Siphonaria diemenensis*, and the black nerite, *Nerita atramentosa*) (Figure 2, Table 1). Although these species provide only a snapshot of the diversity of connectivity patterns expected across our study region, they encompass both active dispersal strategies (bottlenose dolphins and common dolphins, herein ‘*active dispersers*’), and advection-driven larval dispersal strategies (nerites, limpets, and snapper, herein ‘*larval*’ dispersers). Although snapper has a relatively long pelagic larval stage, our categorisation is an oversimplification because the species is also known to move during juvenile and/or adult stage (for details see Bertram et al. 2022, Bertram et al. 2023). Our study taxa include key species for MPA design in South Australia because of their priority status in conservation management plans and fishery stock assessments.

**Figure 2.**
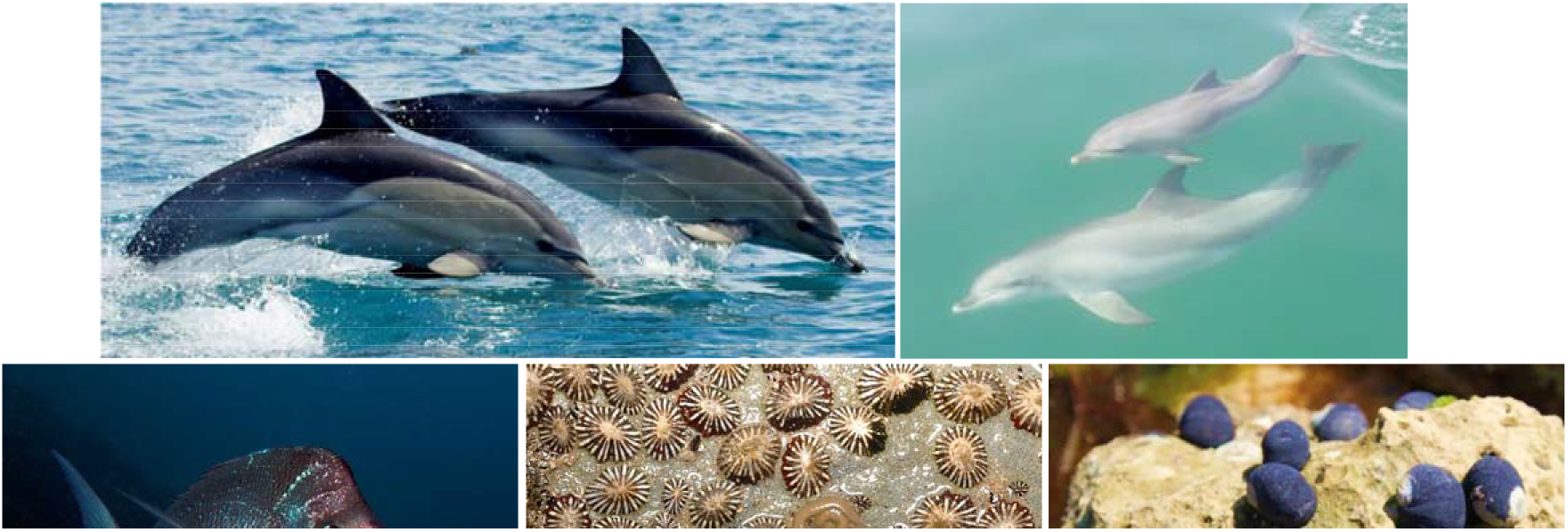
Study species, left to right. Top: *Delphinus delphis* (common dolphin), *Tursiops aduncus* (Indo-Pacific bottlenose dolphin). Bottom: *Chrysophrys auratus* (Australasian snapper), *Siphonaria diemenensis* (Van Diemen’s siphon limpet), *Nerita atramentosa* (black nerite snail). Images by authors or used with permission.

**Table 1.** Sampling information and population genetic differentiation for five species included in the meta-analysis of South Australian Marine Parks network connectivity.

| Species | Taxon class | Data source | Data type | N sites | N indivs | Dispersal | $F_{ST}$ range (& mean) |
| --- | --- | --- | --- | --- | --- | --- | --- |
| *Delphinus delphis* (common dolphin) | Mammalia (mammals) | Barceló et al. (2021); Barceló et al. (2022) | SNP | 5 | 126 | Lifelong, active, potentially year-round | 0.013-0.043 (0.028) |
| *Tursiops aduncus* (Indo-Pacific bottlenose dolphin) | Mammalia (mammals) | Pratt et al. (2022) | SNP | 10 | 117 | Lifelong, active, potentially year-round | 0.035-0.146 (0.090) |
| *Chrysophrys auratus* (Australasian snapper) | Actinopterygii (ray-finned fishes) | Bertram et al. (2022); Bertram et al. (2023) | SNP | 7 | 270 | Larval dispersal up to 30 days, subsequent sub-adult dispersal, decrease in adulthood | 0.007-0.034 (0.021) |
| *Siphonaria diemenensis* (Van Diemen’s siphon limpet) | Gastropoda (snails) | Teske et al. (2016); Teske et al. (2017) | Microsat | 7 | 280 | Larval dispersal 1-2 months | 0.009-0.046 (0.027) |
| *Nerita atramentosa* (Black nerite) | Gastropoda (snails) | Teske et al. (2015); Teske et al. (2017) | Microsat | 9 | 373 | Larval dispersal ~4 months | 0.005-0.013 (0.009) |

SNP = single nucleotide polymorphism (genomic marker); Microsat = microsatellite (genetic marker); N sites = number of South Australian localities sampled; N indivs = total number of individuals; $F_{ST}$ = fixation index representing pairwise genetic differentiation among localities; the scale ranges from 0 (no genetic differentiation) to 1 (complete genetic differentiation).

Based on the localities described in the original publications, we defined sixteen study sites across South Australia (Figure 1), with coverage in 11 of 19 MPA General Use Zones. Since sampling was not originally tailored to a multispecies approach (a limitation common to other datasets worldwide), our approach was limited by incomplete overlap. We chose to retain all unique intra-species sampling localities within the study region (Supplementary Figure 1) but aggregated adjacent inter-species localities into geographically averaged ‘nodes’ to allow for combined analyses, which were labelled according to MPAs in proximity. This strategy was chosen under the assumption that the factors influencing connectivity will be similar across short distances; however, if abrupt biogeographic changes were to occur within these nodes, then inter-species differences would be artificially high. For some species, genetic data were available from outside of South Australia, but the major differences in interstate sampling ranges could also introduce taxonomic bias. We therefore excluded these localities, with the exception of Portland (Locality 16), which is located in the state of Victoria and was sampled for four of the five species, as it was expected to provide data relevant to the nearby, but sparsely sampled, Lower South East bioregion.

### 2.2 Spatial analyses of connectivity across and within the marine parks network

Genetic information was used to quantify population differentiation for each species on a locality-specific basis, as well as pairwise among sampling sites (locality-specific *F*_ST_ and pairwise *F*_ST_, respectively). Locality-specific *F*_ST_ estimates the uniqueness of the ancestry at each locality, relative to the broader dataset or metapopulation (Weir and Hill 2002). This statistic was calculated independently for each study species using the *betas* function in HIERFSTAT 0.5-10 (Goudet 2005) using R (R Core Team 2019), which can be applied to both SNP and microsatellite data. In contrast, pairwise *F*_ST_ estimates differentiation between pairs of localities resulting from population structure (Weir and Hill 2002), and is therefore useful for exploring patterns of divergence across networks. Pairwise *F*_ST_ was calculated using EDENetworks (Kivelä et al. (2015), further described below. A number of indices of differentiation have been proposed as alternatives to *F*_ST_ for markers with high mutation rates, such as microsatellites (e.g., *G’*_ST_, *D* and other indices). For the two microsatellite datasets used here (nerites and limpets), both the overall pattern of very low differentiation across the entire sampled region and the number of pairwise comparisons that were significant were very similar using either *F*_ST_, *G’*_ST_, or *D* (details and statistical comparisons in Teske et al. 2015; Teske et al. 2016). This indicates that *F*_ST_ is an adequate index for our comparative study, which includes microsatellite and SNP datasets. We point to the literature (e.g., Whitlock 2011) for additional information about the choice of an index of differentiation.

To assess the extent to which patterns of connectivity were either shared or unique among the five species, we used a ‘genogeographic’ clustering method (Arranz et al. 2022) to capture relationships between genetic variation and geographic distance along the South Australian coastline for each species. Linearised coastline distances represented the shortest route between a starting point (here, the westernmost locality Nuyts Archipelago Penong), and all other sampling localities while following the coastline. This measure could reflect migration paths of species with nearshore habitat preferences. Linearised coastline distances were first calculated in ArcMap (ESRI 2011) by using the vertex *snap* function to snap sampling coordinates to the nearest segment of the Australian Shoreline layer (Geoscience Australia), and calculating the length of all segments between the starting point and each sampling site. Then, using R code adapted from Arranz et al. (2022), linearised coastline distances were plotted against locality-specific *F*_ST_ values. Curves were fitted to the data points using maximum likelihoods to characterise each species’ spatial trends, and were represented as colour maps depicting variation in genetic divergence along the coast. Fitted curves were scaled and centred, then clustered across species to identify similarities in spatial patterns. Parametric bootstrapping (1000 replicates) of species clustering was used to find the best species clusters, and to assess statistical significance of joins and splits.

### 2.3 Generation and analysis of connectivity networks using a graph theory approach

We used a network approach to summarise patterns of connectivity structure for (a) a combined “*all species*” dataset and (b) for divergent species groupings identified by the genogeographic clustering analysis. First, raw genotype files (SNPs and microsatellites) were imported into EDENetworks to build population-based networks for each species. In each network, nodes (connection points) corresponded to sampling localities, while edges (links among nodes) were calculated as pairwise *F*_ST_ values (Reynolds distance, Reynolds et al. 1983). We exported the resulting distance matrices, before scaling, centring, and averaging species values for each combined network. The distance matrices were then re-imported into EDENetworks for analysis of the combined networks. We derived thresholded networks; a threshold being the maximum pairwise distance considered as providing an effective link between nodes, with all links of larger distances therefore removed. We chose the maximum threshold below percolation, that is, the point at which a connected network would fragment into smaller components. Given the generally high dispersal ability of the species included in this study, the chosen threshold is likely lower than real-world thresholds for these species. However, these thresholds provide an overview of the strongest and weakest pathways in each network, clarifying which localities are likely to become disconnected if overall connectivity is reduced, and which may act as pathways between less connected regions. We calculated network characteristics and node values for each of the thresholded networks and produced circle plots using the R package CIRCLIZE (Gu et al. 2014).

### 2.4 Spatial and oceanographic connectivity models

To assess the relative concordance of spatial distance and oceanographic factors with genetic differentiation across the network, we tested correlations between pairwise *F*_ST_ and explanatory variables of interest (direct waterway distances, coastline distances, latitudinal distances, and ocean advection connectivity estimates), all calculated as pairwise values among all localities. Direct waterway distances refer to the shortest route between each pair of sites without crossing the land, and were calculated using the *viamaris* function in MELFUR (https://github.com/pygmyperch/melfuR). Coastline distances refer to the shortest route between each pair of sites while travelling along the coastline (converted from linearised coastline distances, described in section 2.2). Latitudinal distance was calculated as the difference (in decimal degrees) in latitude between locality pairs, which was included since it could have indirect effects on marine dispersal potential (Álvarez-Noriega et al. 2020).

Pairwise advection connectivity between localities was estimated using the Connectivity Modelling System 1.1 (Paris et al. 2013) to integrate the Ocean General Circulation Model for the Earth Simulator 2 (OFES2; Sasaki et al. 2020). We used a resolution of 0.5° of the 2D velocity fields (eastward and northward) at 5 m depth, from 1994 to 2014. The resulting connectivity matrices show how many particles (e.g., larvae) released from each locality are expected to settle within the same or another sampling locality. We created four matrices representing each season. For each model, we released 1,000 particles per sampling site per day during the three-month seasonal period (a total of 1,800,000 particles per site per model). The particles were advected for at least 30 days before they could settle, and up to 150 days before they were considered dead, approximating the range of larval durations of the three study species with planktonic larvae (*C. auratus*, *N. atramentosa*, and *S. diemenensis*, Table 1). The particle locations were recorded every 3 hours, whereupon it was determined whether they had settled or died. A particle was considered settled when, for the first time, its location intersected within the 1° semicircle surrounding a release site. Because values between localities differed by several orders, estimates were corrected to their natural logarithm. Following the methods of Teske et al. (2015) implemented in our study region, we subjected simulation results to a stepping-stone model of dispersal, by which pairwise advection connectivity was defined as the total number of migrants between each pair of localities after four successive reproductive cycles.

We used redundancy analyses (RDAs) in VEGAN (Oksanen et al. 2019) to test relationships between these variables and the genetic differentiation between localities (pairwise *F*_ST_, as used in the network analyses). Since RDAs do not handle missing data, and not all localities were sampled for all species, we first used a principal component analysis of incomplete data (INDAPCA, Podani et al. 2021) to find the first significant principal components (PCs) of genetic variation. Separate RDAs were then performed for respective environmental variables, where environment acted as an explanatory variable, and genetic PCs acted as a response variable. As with the network analyses, this was performed for a combined *all species* dataset. We repeated analyses for the best -performing models using species subclusters identified by the genogeographic analysis. ANOVAs (function ‘*anova.cca*’) were used to assess the significance of each model with 1000 permutations.

### 2.5 Conservation prioritisation of MPAs

To translate our findings into practical conservation guidance, we applied the genetic connectivity results to spatial prioritization analyses using the conservation planning tool Marxan (Ball et al. 2009). Our aims were to identify priority conservation areas for connectivity within the existing SARSMPA, and to compare prioritisation models based respectively on genetic and spatial connectivity values. The planning units comprised the 16 MPA sample site nodes, with conservation values based on the network ‘betweenness connectivity’ measures for each node obtained from a) independent genetic connectivity networks for each of the five species (five “species” values), b) combined connectivity networks of *larval dispersers* (common dolphins and bottlenose dolphins) and *active dispersers* (snapper, limpets, and nerites) respectively (two “species” values), and c) a connectivity network based on direct waterway distances (a single “species” value). Since conservation targets were hypothetical, we tested iterations with a range of proportional targets (0.2, 0.3, 0.5, 0.7), equal among conservation values in each run of 1,000,000 iterations. Since additional biodiversity features and planning unit ‘costs’ were already integrated into the initial design of the South Australian MPA network, these were not included.

## 3. Results

### 3.1 Spatial connectivity along the network for a range of taxa

Based on the available species’ datasets, we found reasonably high connectivity across the SARSMPA (Figure 3). The lowest population genetic structure (and therefore highest connectivity) was observed for *larval dispersers*; this was reflected by comparatively low pairwise *F*_ST_ values (Table 1). This was most pronounced in nerite snails (average *F*_ST_ = 0.009), followed by snapper (0.021), limpets (0.027), common dolphins (0.028), and bottlenose dolphins (0.090).

**Figure 3.**
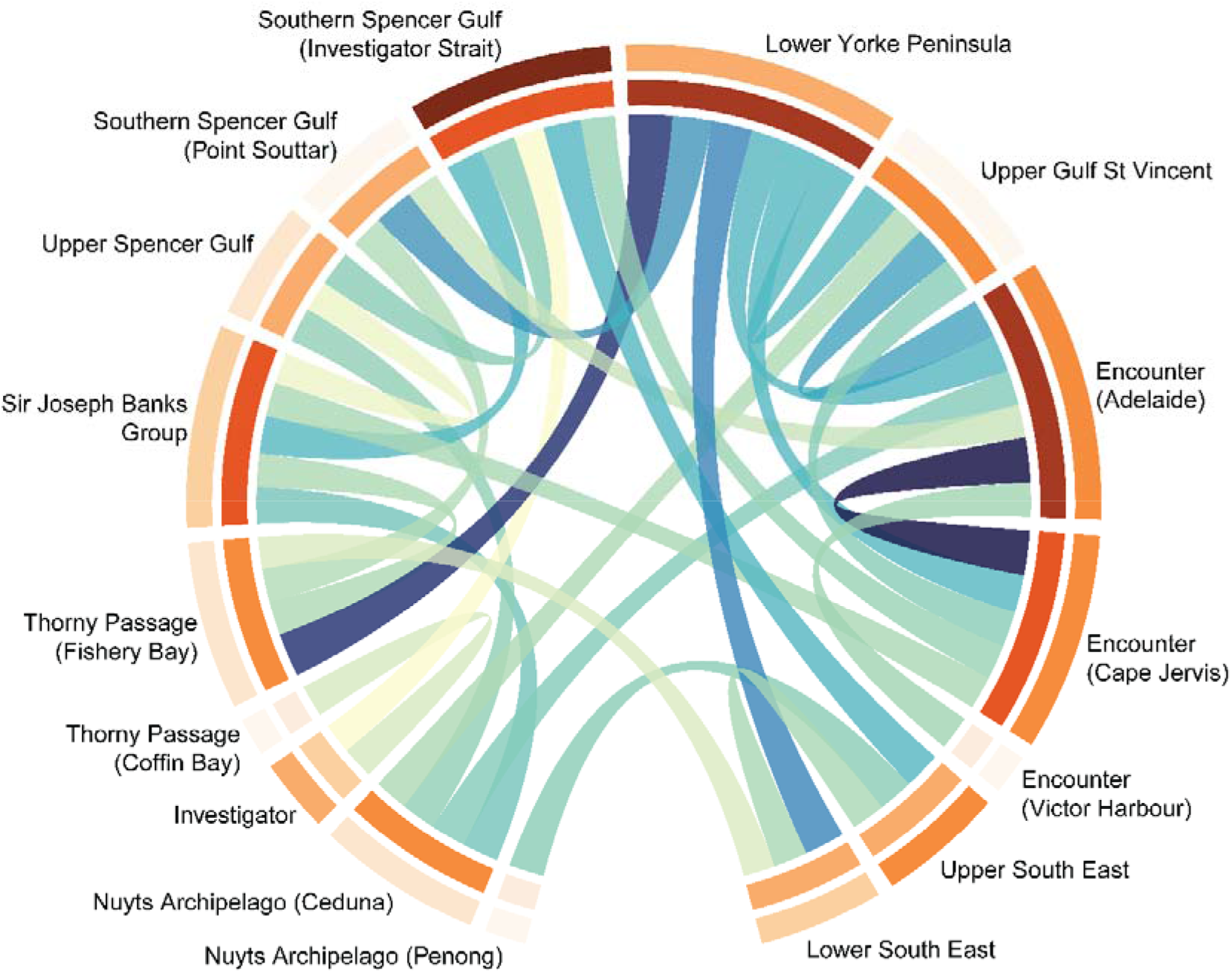
Average connectivity network for all species combined (*Delphinus delphis*, *Tursiops aduncus*, *Chrysophrys auratus*, *Siphonaria diemenensis*, and *Nerita atramentosa*) among sampled marine parks, showing the maximum distance threshold for a fully connected graph. For edges (links in the network), relative pairwise connectivity among nodes is indicated by the degree of shading of the links, with the lightest (yellow) indicating lowest connectivity, and the darkest (navy) indicating the highest connectivity. On surrounding tracks, relative node values are represented by the degree of shading of the orange tracks, with lightest orange indicating the lowest values, and darkest orange indicating the highest values. The outer track represents betweenness centrality (i.e., the node’s importance in forming a pathway between less connected subclusters), while the inner track represents node degree (the total number of links at maximum distance threshold).

Spatial patterns of connectivity were most divergent between *larval dispersers* and *active dispersers*. Genogeographic clustering (Figure 4a; Supplementary Figure 2), based on locality-specific *F*_ST_ values, produced dendrogram groupings with common dolphins and bottlenose dolphins together on one branch, and nerite snails, limpets, and snapper on the other. When average values of each cluster were mapped along the South Australian coastline (Figure 4b), the active-dispersing dolphin species appeared to have higher connectivity in open stretches of coast compared to gulf waters and embayments. In contrast, *larval dispersers* tended to have high connectivity in the centre of the sampling range, especially around the southern reaches of the gulfs. However, despite these trends, associations between no pairs of species were statistically significant (*p* = 0.482-0.597), and siphon limpets were relative outliers within the larval-dispersing cluster. This suggests that despite some shared trends, there also remains a substantial degree of idiosyncrasy in connectivity patterns of individual species.

**Figure 4.**
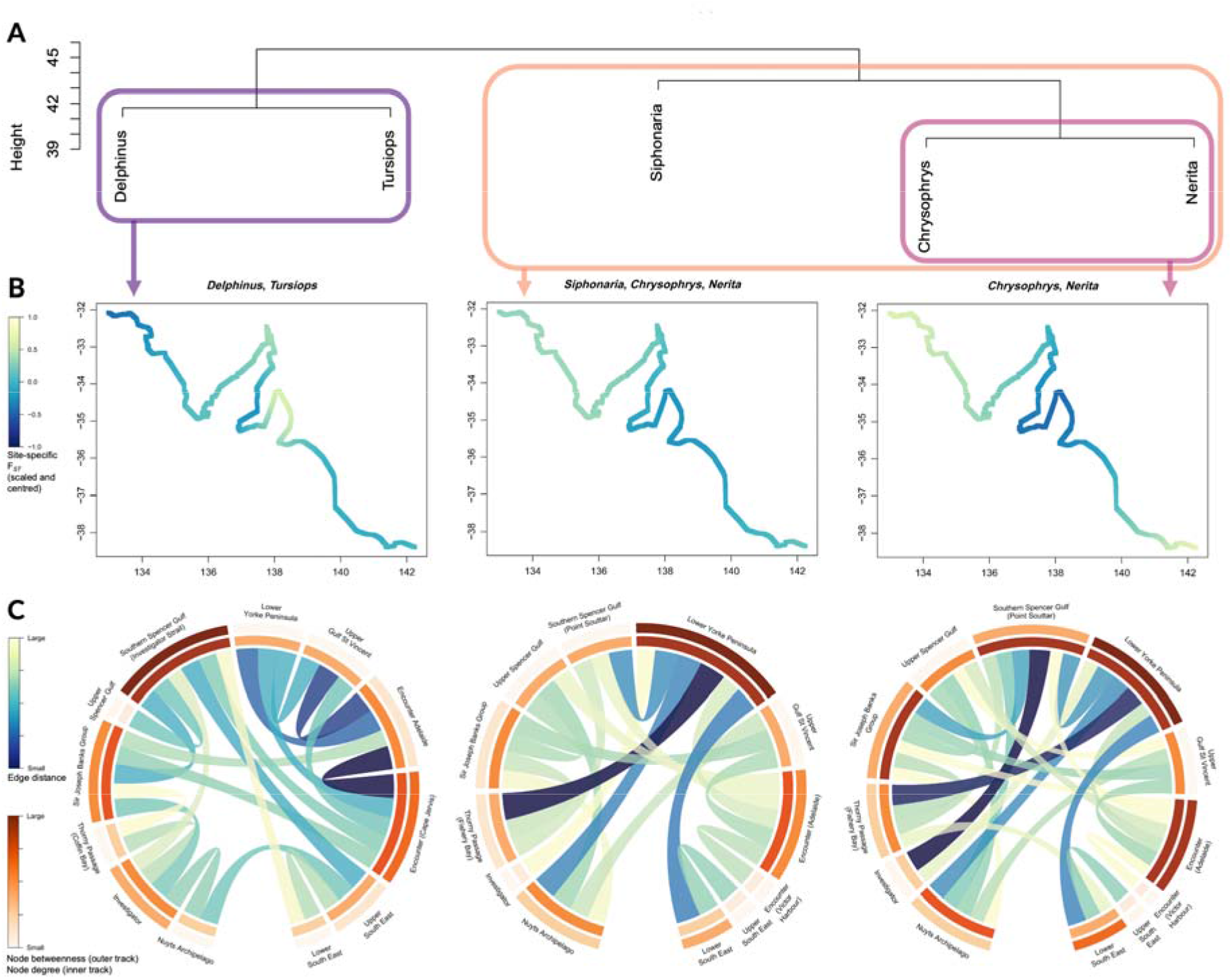
(A) Species clustering based on spatial variation in site-specific genetic differentiation (*F*_ST_); (B) average site- specific genetic differentiation (*F*_ST_) for each species cluster in relation to position along coastline, with lighter yellow indicating regions of greater differentiation/uniqueness; and (C) networks of average connectivity among sampled marine parks per species cluster, showing minimum distance threshold for a fully connected graph. For edges (links in the network), relative pairwise connectivity among nodes is indicated by the degree of shading of the links, with the lightest (yellow) indicating lowest connectivity, and darkest (navy) indicating the highest connectivity. On surrounding tracks, relative node values are represented by the degree of shading of the orange tracks, with lightest orange indicating the lowest values, and darkest orange indicating the highest values. The outer track represents betweenness centrality (i.e., the node’s importance in forming a pathway between less connected subclusters), while the inner track represents node degree (the total number of links at maximum distance threshold).

### 3.2 Network characteristics

At the maximum threshold below percolation (fragmentation of the network), the *all species* dataset network contained 16 nodes (localities), 28 edges (links among localities), and an average node degree (number of connections per node) of 3.5 (Figure 3, Supplementary Tables 1 & 2). The clustering coefficient, reflecting substructure in the network, was 0.29 (where 0 = no substructure and 1 = total substructure). The greatest node degrees were observed for the Lower Yorke Peninsula and Encounter (Adelaide) localities, which each had six strong connections to other nodes. Nodes with only a single strong connection (a node degree of 1) were Nuyts Archipelago (Penong), Thorny Passage (Coffin Bay), and Encounter (Victor Harbor). Also of interest were nodes with high betweenness centrality values, which included Southern Spencer Gulf (Investigator Strait) (38.2), Encounter (Cape Jervis) (19.4), Encounter (Adelaide) (18.9), and Upper South East (17.5).

We also conducted network analyses for active and passive dispersal clusters identified by the genogeographic clustering (Figure 4c). These networks each contained only 12 localities (nodes) due to less sampling coverage within species subsets. For *active dispersers* (dolphins), a thresholded network was produced with 17 edges, an average node degree of 2.83, and a clustering coefficient of 0.31. Southern Spencer Gulf (Investigator Strait) had the greatest node degree of 6, while the nearby Upper Spencer Gulf had the lowest node degree of 1. Since limpets were relatively outlying among *larval dispersers*, we analysed subsets with and without their inclusion (snapper, nerite snails, and limpets; versus snapper and nerites only). The *full larval group* had fewer connecting edges than the *reduced larval group* (18 vs 24), a lower average node degree (3 vs 4), and less clustering (0.21 vs 0.47). Both networks had maximum node degrees of 6. Lower Yorke Peninsula had the highest node degree and betweenness centrality in both full and reduced larval groups. This was the only locality with six connections in the full group, however in the reduced group, three other localities also had node degrees of 6, namely Encounter (Adelaide), Southern Spencer Gulf (Point Souttar), and Sir Joseph Banks Group. Lowest node degrees for both groups were in Encounter (Victor Harbor) and the Upper South East, as well as Investigator in the full larval group.

### 3.3 Spatial and oceanographic relationships with empirical connectivity

We found that genetic connectivity was significantly correlated with both spatial distance and advection connectivity estimates, with differing extents depending on the species subset. For *all species*, direct waterway distance was the best predictor of population connectivity, whereby connectivity declined, and population structure increased, with increasing distances. This relationship accounted for 18.5% of variation of genetic PC1, (*p* <0.001; Figure 5 (upper), Supplementary Table 3). Spring advection was also a relatively good model (associated with 14.9% variation, *p* <0.001, Figure 5 (lower)). While considering these distance and advection variables together could potentially improve predictive power, we also found high autocorrelation between the two (−0.58, Supplementary Figure 3), indicating that combining them could artificially inflate the strength of the models. When considering species by their dispersal clusters, we found that both spatial and advection models were most effective at predicting genetic connectivity in the *active dispersers* (dolphins), compared to any other species cluster. The strongest associations were with the spring advection connectivity model, associated with 36.8% of observed genetic variation (*p* <0.001), followed by direct waterway distances, associated with 33.3% of variation (*p* <0.001). For the *larval dispersers*, direct waterway distance was the only variable significantly associated with genetic connectivity (8.1% of variation, *p* = 0.043). The advection model with the greatest explanatory power was autumn (1.5%, *p* = 0.386), however this was also the variable most highly correlated with distance, which could explain the stronger effect.

**Figure 5.**
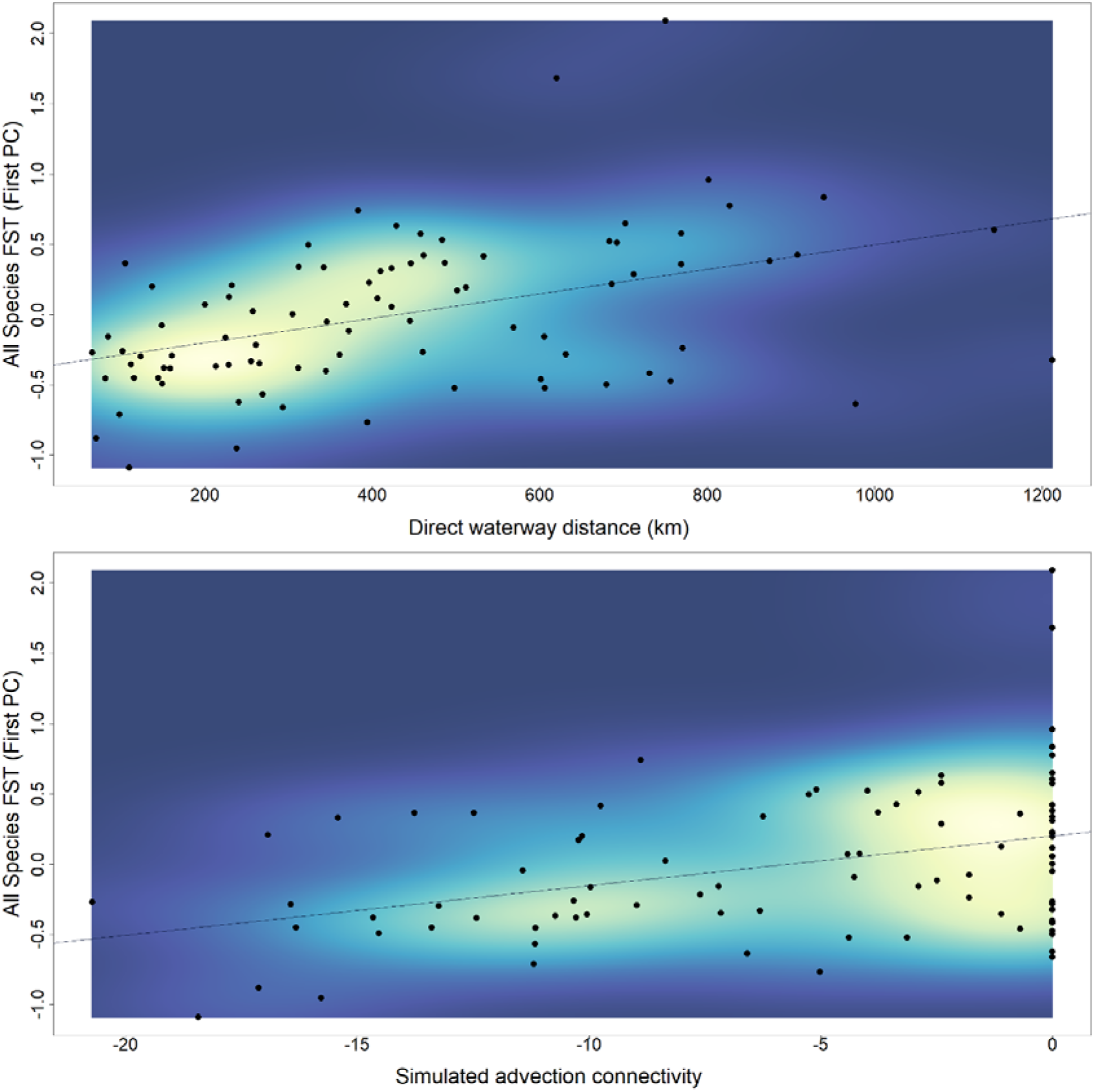
Relationships between multispecies population differentiation and possible explanatory values, with scatterplots showing the line of best fit under a linear regression. Population differentiation is based on first principal component of pairwise *F*_ST_ among South Australian localities for integrated genetic data from the common dolphin (*Delphinus delphis*), bottlenose dolphin (*Tursiops aduncus*), snapper (*Chrysophrys auratus*), siphon limpet (*Siphonaria diemenensis*), and nerite (*Nerita atramentosa*). Top: against direct waterway distances (km), r^2^ = 0.185, *p* <0.001. Bottom: against the best fitting advection connectivity model (spring, steppingstone), r^2^ = 0.149, *p* <0.001.

### 3.4 Spatial conservation prioritisation

Across all Marxan scenarios (different connectivity measures and proportional targets), the nodes most often included in the ‘best’ solution for betweenness connectivity prioritisation were Southern Spencer Gulf (Point Souttar), Southern Spencer Gulf (Investigator strait), and the Upper South East (Supplementary Table 4). MPAs that were never prioritised included Thorny Passage (both Coffin Bay and Fishery Bay), Upper Spencer Gulf, Upper Gulf St Vincent, and Encounter (Victor Harbor). There was a significant, although modest, correlation between selection frequencies (Figure 6, Supplementary Table 5) among genetic and spatial prioritisation run (*p* = 0.0006, 0.226). Notably, several MPAs were consistently selected in some connectivity scenarios, but not in others. An interesting example was that Investigator was the most consistently prioritised node in spatial connectivity scenarios, but was never prioritised in any genetic connectivity scenario, demonstrating the possibility for prioritisation discrepancies. A larger number of MPAs were typically priorities in best solution for the five species genetic scenario, however, this is likely an artifact of the inclusion of a larger number of ‘conservation features’ requiring priority, and can be balanced by comparing lower proportional targets of this scenario with higher proportional targets of other scenarios.

**Figure 6.**
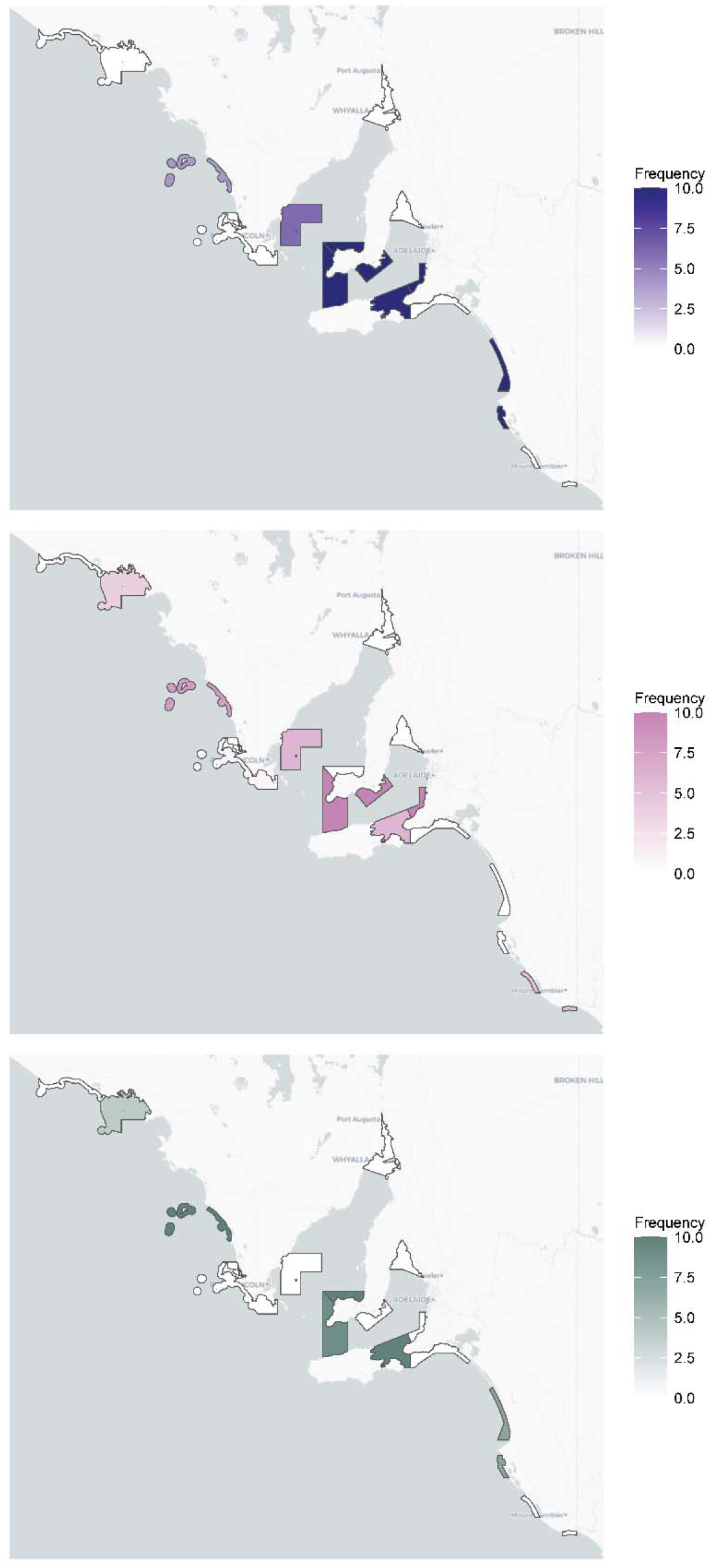
Selection frequency of planning units (marine park segments) using Marxan under 1,000,000 iterations, where conservation feature values are based on node ‘betweenness’ from a) independent genetic connectivity networks of all five species, b) combined connectivity networks of “larval dispersers” and “active dispersers”, and c) a connectivity network based on direct waterway distances.

## 4. Discussion

### 4.1 An approach for integrating genetic connectivity into MPA network planning

We implemented an integrative approach to incorporate population connectivity as a factor in the planning and assessment of MPAs. Our case study focused on existing management zones within a network of MPAs, relying on genetic data from multiple species with shared or nearby sampling localities across the region. The approach requires neutral genetic or genomic data (i.e., of no adaptive relevance) to generate metrics of pairwise and site-specific genetic differentiation. By treating the sampled MPA zones as nodes in a network, we were able to evaluate their contribution to regional connectivity. This evaluation encompassed measures of connectivity for all represented species, as well as for species clusters which were found to have more similar connectivity patterns. In this case, patterns were more closely shared among species with similar dispersal strategies, emphasising the importance of selecting taxa that are representative of the diversity of various life history traits. We demonstrated the use of these results in spatial conservation planning, using Marxan to identify priority areas while considering hypothetical connectivity targets. Furthermore, we evaluated proxy connectivity values, including geographic distances and advection dispersal models. Among these potentially predictive metrics, we found that direct waterway distances exhibited the highest correlation with multispecies connectivity in the MPA network, validating their previous use in planning when genetic connectivity data was lacking. Nevertheless, our study revealed notable inconsistencies between the prioritisation models derived from genetic connectivity networks and those from geographic distance networks. These discrepancies emphasise the value of incorporating genetic data in spatial conservation planning, even when considering other representative measures.

### 4.2 Connectivity within the MPA network

High effective network connectivity across all taxa was consistent with general expectations for long-range dispersers (Waples 1998), which could describe all included species. However, the Southern Spencer Gulf (Investigator Strait), Southern Spencer (Point Souttar), and the Lower Yorke Peninsula were the most important hubs of connectivity across *all species* networks. These localities not only had the greatest number of strong connections with other sites, but were also ranked highly for betweenness centrality, indicating their potential for gene flow relay between more disconnected areas (Kivelä et al. 2015). The two localities provided links to the less connected gulf waters, and had relatively strong connectivity with distant MPAs, including the westernmost Nuyts Archipelago (for *larval dispersers*) and easternmost Upper South East and Lower South East (for all dispersal groups). Since the Investigator Strait represents a transition zone between gulf waters and pelagic waters, this area may also represent high connectivity between inshore and offshore communities (Scientific Working Group 2011). The existence of two MPAs in this area, including five sanctuary zones, is therefore positive for the maintenance of ecological connectivity across the SARSMPA network. A focus on monitoring and compliance should be a priority in this part of the network to maximise the protection of representative habitats and species.

High connectivity values may be influenced by hub localities’ orientation at the centre of a sampling range; however, this did not appear to apply to upper gulf waters, despite their relative longitudinal centrality. Lower average connectivity of the gulfs compared to surrounding localities was consistent with influences of front formations at the gulf entrances, which is thought to allow accumulation of high densities of fish larvae in Investigator Strait during warmer months, but limits passive dispersal into the above gulf regions (Bruce and Short 1990, Fowler et al. 2000). Gulf waters and embayments have also been associated with higher site-fidelity and residency in dolphins (Bilgmann et al. 2007, Möller et al. 2007, Fruet et al. 2014, Passadore et al. 2018a). This may be a contributing factor in the particularly low connectivity of the *active dispersers* group between gulfs and the surrounding stretches of coastline. The stronger network clustering of *active dispersers* was also consistent with high site fidelity in the gulf waters. At network thresholds above percolation (i.e., if the weakest links in the network were to be removed), network breakdown would likely first occur between the two gulf-associated subclusters. Given the prior lack of data about multispecies connectivity patterns, this information provides a baseline for understanding the structure of South Australian marine park connectivity and for informing ongoing monitoring practices.

### 4.3 Environmental correlations and potential drivers of connectivity

We hypothesised that patterns of genetic connectivity would be associated with variations in geographic distance and oceanographic circulation, which was supported by strong associations with direct waterway distance and spring advection connectivity for the *all-species* dataset. However, at this level, advection models did not improve predictions over distance alone. Moreover, we found that the best explanatory variables were not shared among life history subclusters (*active dispersers* versus *larval dispersers*). This suggests that idiosyncrasies among these groups (and potentially among comprised species) could limit the generalisability of oceanographic modelling for predictions in multispecies networks, at least within similar data constraints, as seen here. Since advection models are being increasingly applied to connectivity estimates based on general dispersal traits rather than empirical data (e.g. Jonsson et al. 2016, Bray et al. 2017, Roberts et al. 2021), ongoing calibration with observational data may be required to ensure accuracy (Faillettaz et al. 2018).

Unexpectedly, we found that advection connectivity better predicted connectivity of *active dispersers* than *larval dispersers*. Given that advection models were approximated from life history considerations of the snapper, limpet, and nerite, we hypothesised that advection models for these species might outperform predictions based on distance alone. Their underperformance for *larval dispersers* could potentially relate to the lack of species-specificity of the models, or might instead be attributed to factors such as habitat suitability, temporal variation in recruitment, inappropriate spatial scale, or to unknown barriers in the intervening matrix (Hedgecock 1994, Banks et al. 2007, Teske et al. 2015). Equally interesting was the very strong explanatory power of the spring advection model for the *active dispersers*’ connectivity, despite its design for larval predictions. However, this was not a novel finding: strong associations have previously been found between oceanography and dolphins’ population divergence in southern Australia, with possible relevance to hydrological adaptation and feeding specialisations (Bilgmann et al. 2007, Barceló et al. 2022, Pratt et al. 2022). It is plausible that similar factors may be contributing to structure across current-driven habitat gradients in the present study. Despite ongoing questions, a useful takeaway from the association analyses was the overall utility of geographic distance as a metric for estimating broad patterns of connectivity across species of varying life histories. While there is room for improving the predictive power of both current and future modelling, these results help to affirm established planning and evaluation practices in the SARSMPA, which have used spatial measures as a surrogate for connectivity metrics for spatial prioritisation within the network (DEH 2008). However, it is essential to highlight that ‘best solutions’ from spatial prioritisation models were in several cases mismatched between those reliant on genetic connectivity networks and those based on geographic distances.

Connectivity can promote adaptation and increase resilience to climate change in a number of ways. First, the exchange of genetic variation among populations or metapopulations can help to maintain local reservoirs of standing diversity — the raw material on which natural selection can act (Hendry and Taylor 2004, Nosil et al. 2019). Second, existing climatic variation across species’ habitat ranges can lead to divergent regional adaptations, and some ‘pre-adapted’ genetic variations may spread quickly throughout the network if they become favourable under changing climates (Haldane 1948, Nosil et al. 2019). Lastly, connectivity can simply permit individuals to relocate to more suitable conditions if they have access to the required habitats. Recently, putative genetic adaptations in association with gradients in sea surface temperature across South Australia have been identified in both Indo-Pacific bottlenose dolphins (Pratt et al. 2022) and common dolphins (Barceló et al. 2022). However, since less dispersive dolphins resident to embayment habitats are considered more likely to be impacted by extreme heatwaves (Wild et al. 2019, Barceló et al. 2022), the capacity for gene flow and dispersal is particularly valuable. In the present study, network analyses indicated that the strongest link between dolphin population subclusters was between Encounter (Adelaide) and Southern Spencer Gulf (Investigator Strait). Without marine park zoning, dolphins in these regions are at increased risk from fisheries interactions, habitat degradation, coastal zone development, and recreational activities (Hamer et al. 2008, Passadore et al. 2018b, Barceló et al. 2021). Population depletion at these important nodes could have wide-ranging effects by reducing connectivity and adaptive genetic exchange between gulf systems, and thereby the wider network. Modelling oceanographic changes in future climate scenarios may help to pre-empt changes to connectivity pathways; however, a more immediate priority will be to refine understandings of the associations between oceanographic connectivity models and observed genetic structure.

### 4.4 Limitations and recommendations

This study was designed to capitalise on existing genetic information spanning an MPA network. However, an important limitation was the number of datasets available from this region, which we acknowledge increases the risk of taxonomic or life history-related biases, especially at sites with incomplete overlap of sampling efforts. While multiple analytical steps were taken to reduce possible biases, the distinctness of species-specific connectivity patterns suggests that a broader representation and more intensive sampling would greatly strengthen inferences about region-wide patterns. Our recommendations for future studies therefore include maximising both the number and variety of taxa, and the overlap of sampling localities, ideally with an understanding of whether missing node data are due to previous sampling strategies or genuine absence. If enough funding is attainable, it may be possible to supplement these data with sampling tailored to address the deficiencies identified in a particular region. We also suggest that efforts to enhance collaboration among researchers, curators, and data providers can facilitate the compilation of standardised, multi-species genetic datasets. This will contribute to more robust and inclusive assessments of marine connectivity, and improve the overall effectiveness of conservation efforts within marine protected areas.

Assuming community compliance with marine park regulations, we can suggest that the current distribution of Marine Parks in the SARSMPA is expected to help maintain ecological processes associated with connectivity and climatic resilience. The results of this study will serve as a baseline for ongoing connectivity assessments, which we recommend be extended by including genetic data. While the results support the usefulness of spatial proxies in connectivity planning, there remains great scope to extend connectivity assessments for a more comprehensive understanding of spatial, temporal, and biological influences on linkages within the network. In the face of climate change and increasing anthropogenic pressures, continued integration of biological and physical data will be invaluable to the sustainable management of marine ecosystems.

#### CRediT authorship contribution statement

**Katie Gates:** Formal analysis, Writing - Original Draft. **Jonathan Sandoval-Castillo:** Formal analysis, Investigation, Writing - Review & Editing. **Andrea Barceló, Andrea Bertram, Eleanor Pratt, Peter Teske:** Investigation, Writing - Review & Editing**. Luciano Beheregaray and Luciana Möller**: Conceptualization, Supervision, Project administration, Resources, Investigation, Funding acquisition, Writing - Review & Editing.

## Acknowledgments

This work was funded by the South Australian Department for Environment and Water (DEW) research grant *Monitoring and Evaluation of South Australia marine parks network*. We are grateful to Craig Meakin, Danny Brock, and Simon Bryars for their discussions and feedback.

## Data availability

Data will be made available in *figshare* upon manuscript acceptance.

