## Supplemental Files for "Connecting the dots: applying multispecies connectivity in marine park network planning"

Appendix 1: Supplementary Material

Supplementary Figures

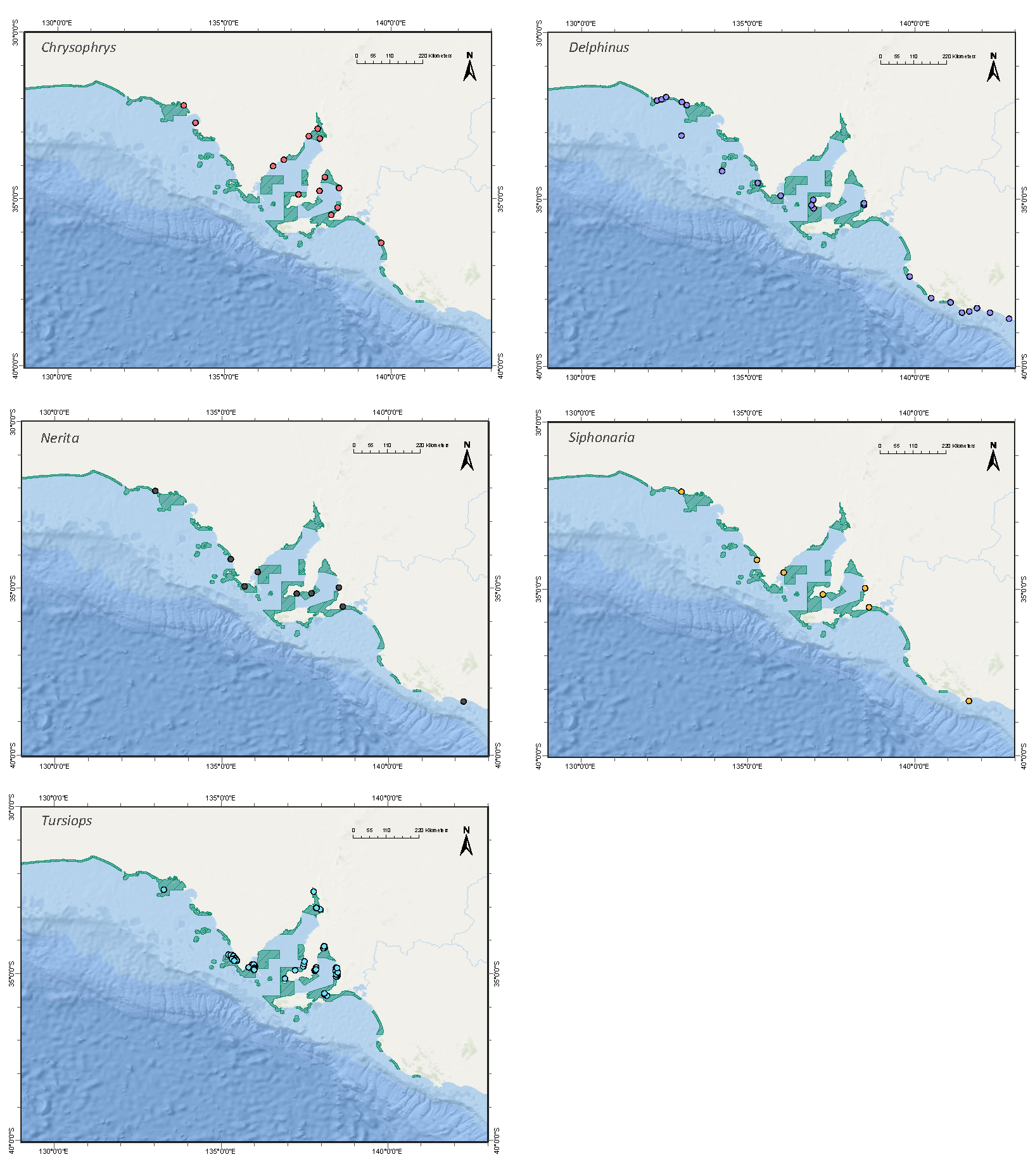

Supplementary Figure 1. Sampling localities, indicated by coloured circles, across the South Australian Representative System of Marine Protected Areas (SARSMPA) for five species, *Chrysophrys auratus*, *Delphinus delphis*, *Nerita atramentosa*, *Siphonaria diemenensis*, and *Tursiops aduncus*. SARSMPA General Managed Use Zones are indicated by green shading.

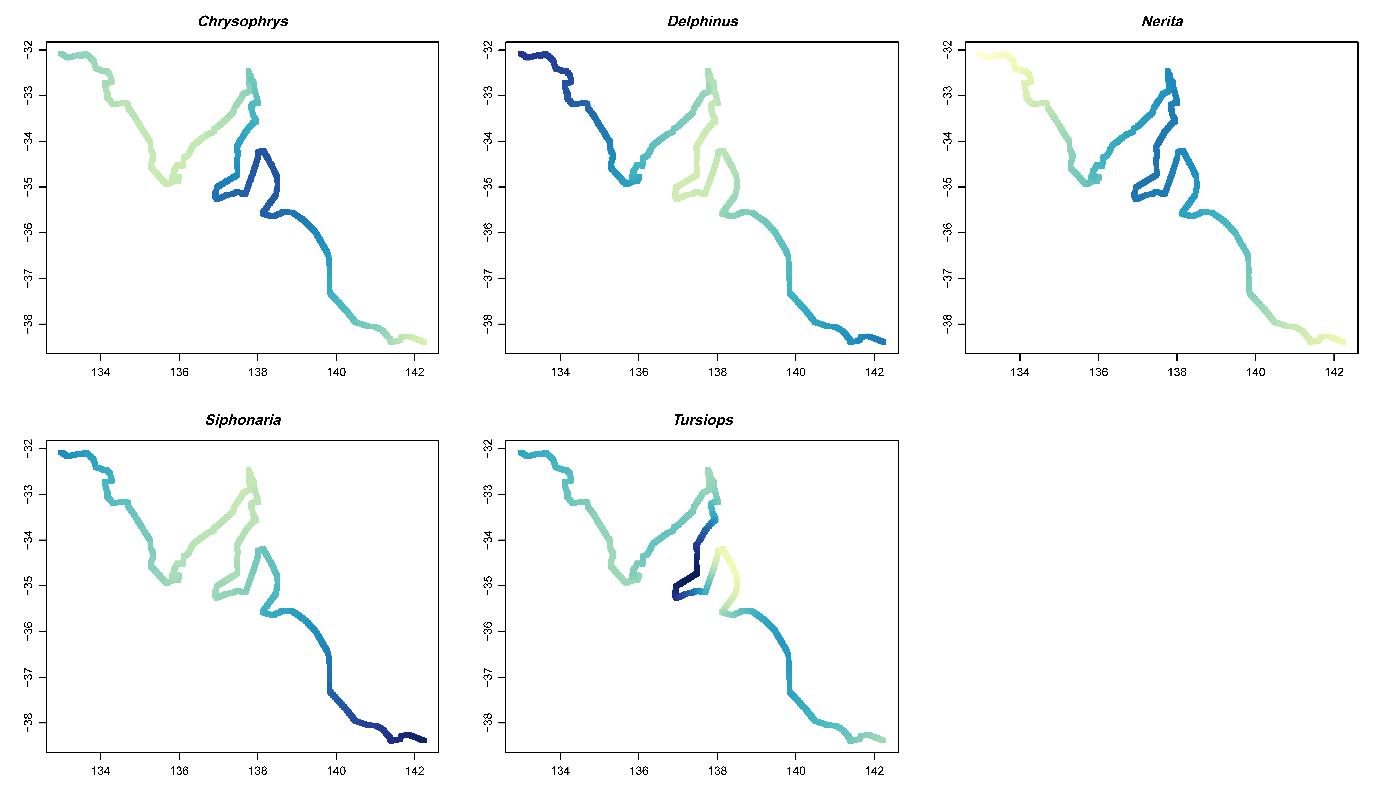

Supplementary Figure 2. Individual genogeographic maps for each species’ average site-specific genetic differentiation (*F*_ST_) for five species, *Chrysophrys auratus*, *Delphinus delphis*, *Nerita atramentosa*, *Siphonaria diemenensis*, and *Tursiops aduncus*, in relation to position along the coastline, with lighter yellow indicating regions of greater differentiation/uniqueness.

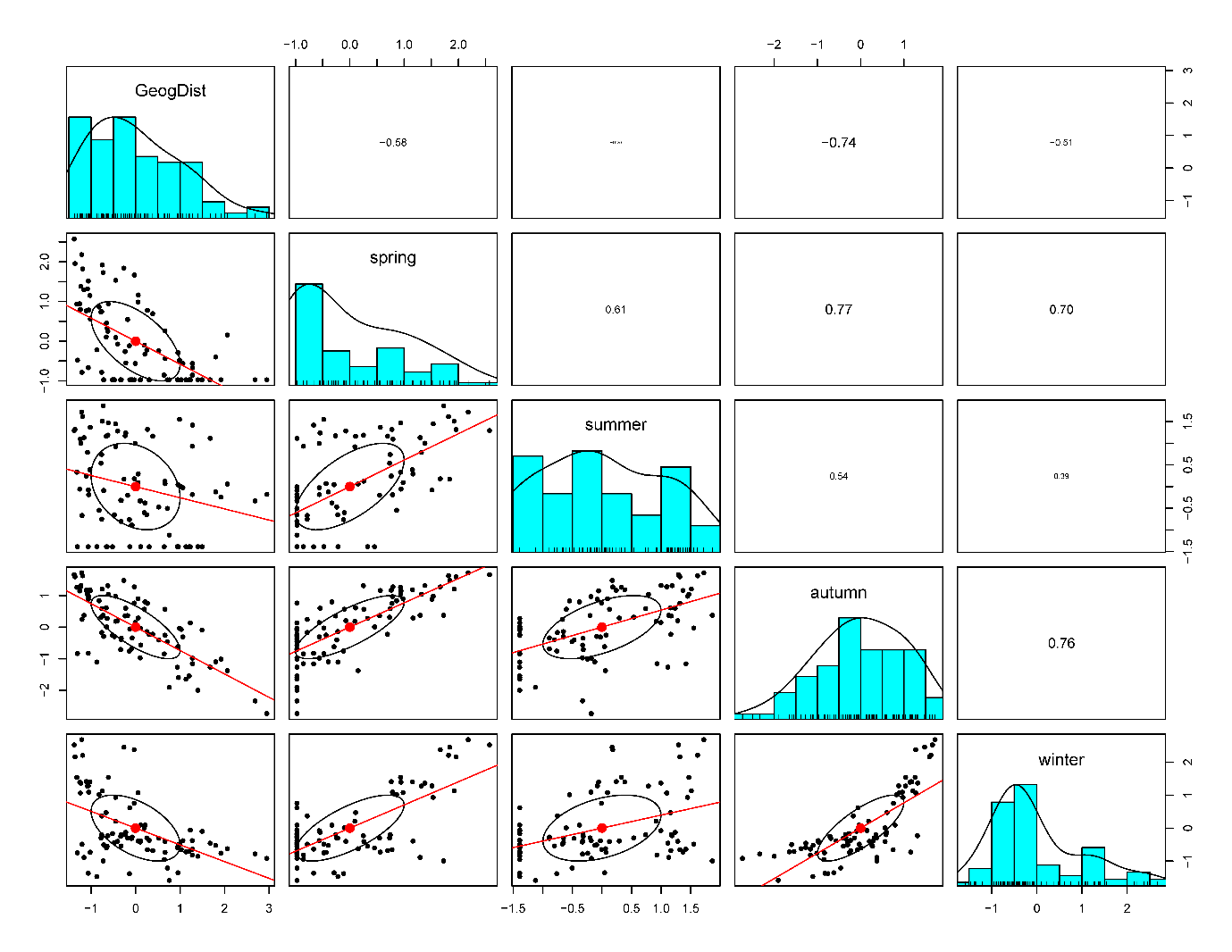
 Supplementary Figure 3. Proportion of autocorrelation among environmental variables used in RDAs. GeogDist = direct waterway distances. Seasons correspond to advection connectivity models calculated for respective three-month periods, based on the Ocean General Circulation Model for the Earth Simulator 2 (OFES2; Sasaki et al. 2020).

Supplementary Tables

Supplementary Table 1. Characteristics of thresholded connectivity networks (main results, Figures 3 & 4), where dmax = maximum network distance (equivalent to percolation threshold), N = number of nodes (localities), E = number of edges (connections in the thresholded networks), <k> = average node degree (average number of connections per node), kmax = maximum node degree (maximum number of connections of any node in network), and <c> = clustering coefficient (scale 0-1).

|  | **All species** | **Active Dispersers** | **Larval Dispersers** | **Reduced Larval** |
| --- | --- | --- | --- | --- |
| **dmax** | 0.323 | 0.288 | 0.314 | 0.443 |
| **N** | 16 | 12 | 12 | 12 |
| **E** | 28 | 17 | 18 | 24 |
| **<k>** | 3.5 | 2.83 | 3 | 4 |
| **kmax** | 6 | 6 | 6 | 6 |
| **<c>** | 0.29 | 0.31 | 0.21 | 0.47 |

Supplementary Table 2. Node data for network analyses across 16 sampling localities in the South Australian Marine Parks network. Node degree = number of connections per node. Betweenness centrality = the number of shortest paths between other nodes which must pass through that node.

| **Locality** | **Node Degree** | | | |  | **Betweenness Centrality** | | | |
| --- | --- | --- | --- | --- | --- | --- | --- | --- | --- |
|  | All species | Active Dispersers | Larval Dispersers | Reduced Larval |  | All species | Active Dispersers | Larval Dispersers | Reduced Larval |
| Nuyts Archipelago Penong | 1 | 2 | NA | NA |  | 0.0 | 0.8 | NA | NA |
| Nuyts Archipelago Ceduna | 4 | NA | 4 | 5 |  | 5.4 | NA | 6.3 | 4.1 |
| Investigator | 2 | 4 | 1 | 2 |  | 14.0 | 8.3 | 0.0 | 0.0 |
| Thorny Passage Coffin Bay | 1 | 2 | NA | NA |  | 0.0 | 0.0 | NA | NA |
| Thorny Passage Fishery Bay | 4 | NA | 3 | 4 |  | 6.5 | NA | 3.0 | 4.6 |
| Sir Joseph Banks Group | 5 | 5 | 4 | 6 |  | 11.6 | 13.0 | 2.8 | 5.8 |
| Upper Spencer Gulf | 3 | 1 | 3 | 4 |  | 2.5 | 0.0 | 1.7 | 0.4 |
| Southern Spencer Gulf Point Souttar | 3 | NA | 3 | 6 |  | 1.0 | NA | 1.0 | 5.9 |
| Southern Spencer Gulf Investigator Strait | 5 | 6 | NA | NA |  | 38.2 | 26.0 | NA | NA |
| Lower Yorke Peninsula | 6 | 3 | 6 | 6 |  | 13.7 | 0.0 | 29.2 | 15.9 |
| Upper Gulf St Vincent | 4 | 3 | 3 | 4 |  | 1.4 | 0.0 | 1.7 | 0.4 |
| Encounter Adelaide | 6 | 4 | 5 | 6 |  | 18.9 | 3.2 | 16.3 | 14.1 |
| Encounter Cape Jervis | 5 | 5 | NA | NA |  | 19.4 | 14.8 | NA | NA |
| Encounter Victor Harbor | 1 | NA | 1 | 1 |  | 0.0 | NA | 0.0 | 0.0 |
| Upper South East | 3 | 3 | 1 | 1 |  | 17.5 | 2.8 | 0.0 | 0.0 |
| Lower South East | 3 | 2 | 2 | 3 |  | 9.0 | 0.0 | 10.0 | 10.0 |

Supplementary Table 3. RDA summary statistics for associations between environmental variables and first PCs of genetic variation for all species, and for species subclusters. For *p*-values, significance codes *p* < 0.001 = ***, *p* < 0.01 = **, *p* < 0.05 = *

| **Environmental variable vs all species connectivity** | | **R^2^** | **Adj R^2^** | ***p*-value** | **F** |
| --- | --- | --- | --- | --- | --- |
| **Distance** | Direct waterway | 0.185 | 0.176 | 0.001 *** | 19.33 |
|  | Coastal | 0.115 | 0.104 | 0.001 *** | 10.99 |
|  | Latitudinal | 0.093 | 0.082 | 0.004 ** | 8.72 |
| **Advection model** | Spring | 0.149 | 0.139 | 0.001 *** | 14.94 |
|  | Summer | 0.033 | 0.021 | 0.102 | 2.88 |
|  | Autumn | 0.113 | 0.103 | 0.002 ** | 10.86 |
|  | Winter | 0.092 | 0.081 | 0.005 ** | 8.62 |
|  | Sum of seasons | 0.103 | 0.092 | 0.002 ** | 9.72 |
| **Environmental variable vs dolphin connectivity** | |  |  |  |  |
| **Distance** | Direct waterway | 0.333 | 0.319 | 0.001 *** | 24.93 |
| **Advection model** | Spring | 0.368 | 0.355 | 0.001 *** | 29.08 |
|  | Summer | 0.266 | 0.251 | 0.001 *** | 18.08 |
|  | Autumn | 0.244 | 0.228 | 0.001 *** | 16.10 |
|  | Winter | 0.172 | 0.156 | 0.004 ** | 10.42 |
| **Environmental variable vs larval disperser connectivity** | |  |  |  |  |
| **Distance** | Direct waterway | 0.081 | 0.062 | 0.043 * | 4.32 |
| **Advection model** | Spring | 0.019 | -0.001 | 0.353 | 0.93 |
|  | Summer | 0.014 | -0.006 | 0.434 | 0.69 |
|  | Autumn | 0.015 | -0.005 | 0.386 | 0.74 |
|  | Winter | 0.036 | 0.016 | 0.191 | 1.81 |

Supplementary Table 4: Marxan ‘best solutions’ for hypothetical connectivity proportional targets (0.2, 0.3, 0.5, 0.7). Planning units comprised the 16 MPA sampling site nodes, with conservation values based on the network ‘betweenness connectivity’ measures for each node obtained from a) independent genetic connectivity networks for each of the five species b) combined connectivity networks of active dispersers and larval dispersers and c) a connectivity network based on direct waterway distances.

|  | Five species independently | | | | Active and larval dispersers | | | | Waterway distances | | | |
| --- | --- | --- | --- | --- | --- | --- | --- | --- | --- | --- | --- | --- |
| Target proportion to conserve | 0.2 | 0.3 | 0.5 | 0.7 | 0.2 | 0.3 | 0.5 | 0.7 | 0.2 | 0.3 | 0.5 | 0.7 |
| NuytsArchipelagoPenong |  |  |  |  |  |  |  |  |  |  |  |  |
| NuytsArchipelagoCeduna |  |  |  |  |  |  |  |  |  |  |  | 1 |
| Investigator |  |  |  |  |  |  |  |  | 1 | 1 | 1 | 1 |
| ThornyPassageCoffinBay |  |  |  |  |  |  |  |  |  |  |  |  |
| ThornyPassageFisheryBay |  |  |  |  |  |  |  |  |  |  |  |  |
| SirJosephBanksGroup |  | 1 | 1 | 1 |  |  |  | 1 |  |  |  |  |
| UpperSpencerGulf |  |  |  |  |  |  |  |  |  |  |  |  |
| SouthernSpencerGulfPointSouttar | 1 | 1 | 1 | 1 |  |  |  |  |  |  | 1 | 1 |
| SouthernSpencerGulfInvestigatorStrait | 1 |  |  | 1 | 1 | 1 | 1 | 1 |  |  |  |  |
| LowerYorkePeninsula |  |  |  | 1 |  | 1 | 1 | 1 |  |  |  |  |
| UpperGulfStVincent |  |  |  |  |  |  |  |  |  |  |  |  |
| EncounterAdelaide |  |  | 1 | 1 | 1 |  | 1 | 1 |  |  |  |  |
| EncounterCapeJervis |  |  |  | 1 |  |  | 1 | 1 |  |  | 1 | 1 |
| EncounterVictorHarbor |  |  |  |  |  |  |  |  |  |  |  |  |
| UpperSouthEast | 1 | 1 | 1 | 1 |  |  |  |  |  | 1 |  | 1 |
| LowerSouthEast |  |  |  |  |  |  |  | 1 |  |  |  |  |

Supplementary table 5: Marxan ‘summed solutions’ for hypothetical connectivity proportional targets (0.2, 0.3, 0.5, 0.7). Planning units comprised the 16 MPA sampling site nodes, with conservation values based on the network ‘betweenness connectivity’ measures for each node obtained from a) independent genetic connectivity networks for each of the five species b) combined connectivity networks of active dispersers and larval dispersers and c) a connectivity network based on direct waterway distances.

|  | Five species independently | | | | | | | Active and larval dispersers | | | | | | | | Waterway distances | | | | | | |
| --- | --- | --- | --- | --- | --- | --- | --- | --- | --- | --- | --- | --- | --- | --- | --- | --- | --- | --- | --- | --- | --- | --- |
| Target proportion to conserve | 0.2 | 0.3 | | 0.5 | | 0.7 | | 0.2 | | 0.3 | | 0.5 | | 0.7 | | 0.2 | | 0.3 | | 0.5 | | 0.7 |
| NuytsArchipelagoPenong |  |  | |  | |  | |  | |  | |  | |  | |  | |  | |  | |  |
| NuytsArchipelagoCeduna |  |  | | 2 | | 4 | |  | |  | |  | |  | |  | | 1 | |  | | 4 |
| Investigator |  |  | | 1 | | 8 | | 1 | | 4 | |  | | 4 | | 9 | | 8 | | 10 | | 10 |
| ThornyPassageCoffinBay |  |  | |  | |  | |  | |  | |  | |  | |  | |  | |  | |  |
| ThornyPassageFisheryBay |  |  | |  | | 1 | |  | |  | |  | |  | |  | |  | |  | |  |
| SirJosephBanksGroup | 1 |  | | 6 | | 6 | | 1 | | 2 | | 10 | | 6 | | 1 | |  | |  | |  |
| UpperSpencerGulf |  |  | |  | |  | |  | |  | |  | |  | |  | |  | |  | |  |
| SouthernSpencerGulfPointSouttar |  |  | |  | |  | | 7 | | 10 | | 10 | | 10 | |  | | 3 | | 10 | | 10 |
| SouthernSpencerGulfInvestigatorStrait | 7 | 10 | | 10 | | 10 | | 5 | |  | |  | | 10 | |  | | 3 | |  | | 9 |
| LowerYorkePeninsula | 3 | 10 | | 10 | | 10 | |  | |  | |  | | 10 | |  | |  | |  | |  |
| UpperGulfStVincent |  |  | |  | |  | |  | |  | |  | |  | |  | |  | |  | |  |
| EncounterAdelaide | 7 |  | | 6 | | 10 | | 5 | | 4 | | 10 | | 10 | |  | |  | |  | |  |
| EncounterCapeJervis | 2 |  | | 3 | | 6 | | 3 | | 4 | |  | | 10 | | 1 | | 4 | | 10 | | 10 |
| EncounterVictorHarbor |  |  | |  | |  | |  | |  | |  | |  | |  | |  | |  | |  |
| UpperSouthEast |  |  | |  | |  | | 8 | | 6 | | 10 | | 10 | |  | | 1 | |  | | 7 |
| LowerSouthEast |  |  | | 2 | | 5 | |  | |  | |  | |  | |  | |  | |  | |  |
